# Genetic ablation of Pth4 disrupts calcium-phosphate balance, bone development, and kidney transcriptome in teleosts

**DOI:** 10.1101/2025.05.13.653078

**Authors:** Luis Méndez-Martínez, Paula Suarez-Bregua, Laura Guerrero-Peña, Elisa Barreiro-Docío, Carolina Costas-Prado, Antonio Cobelo-García, Josep Rotllant

## Abstract

Parathyroid hormone 4 (Pth4) is an evolutionarily conserved member of the PTH family, expressed in hypothalamic neurons and lost in eutherian mammals. In order to elucidate its role in mineral homeostasis and skeletal development, a *pth4* knockout (*pth4*^*KO*^) zebrafish line was generated using CRISPR/Cas9 and transcriptomic profiling was conducted across six key tissues: brain, kidney, intestine, gills, scales, and bone. The results obtained demonstrated that the loss of Pth4 led to pronounced disturbances in calcium and phosphate homeostasis, skeletal deformities, and widespread tissue-specific transcriptional alterations. Notably, dysregulation of mineral regulatory genes— such as *fgf23, phex*, and *slc34a1a* was particularly evident in the kidney, suggesting disruption of the FGF23-Klotho axis. In parallel, differential expression of extracellular matrix genes (*col1a1a, col10a1a, col11a1*) and matrix remodeling enzymes (*mmp9, mmp13a, mmp2*) in bone and scales indicated impaired skeletal remodeling. Together, these findings highlight a pivotal role for Pth4 in the endocrine and local regulation of mineral metabolism and skeletal integrity, expanding our understanding of PTH family functions in vertebrate physiology.

## Introduction

Bone development and mineral homeostasis in vertebrates are complex biological processes. These are regulated by an intricate interplay of hormones, bone cell turnover and regulatory peptides. Of particular significance within this complex network is the parathyroid hormone (PTH) family, which plays a pivotal role in the maintenance of calcium-phosphate balance, bone remodeling and mineralization. In humans, the PTH family comprises three structurally related peptides: PTH, PTH2, and PTHLH. These peptides exert their effects through two G-protein-coupled receptors, PTH1R and PTH2R (John T Potts, 2005). In contrast, teleosts exhibit an expanded PTH family as a result of an additional round of whole-genome duplication, leading to at least six PTH-related peptides: Pth1a, Pth1b, Pth2, Pth3a (also known as Pthlha), Pth3b (also known as Pthlhb) and Pth4 (Glasauer & Neuhauss, 2014; Guerreiro et al., 2007; Suarez-Bregua, Torres-Nuñez, et al., 2017). These peptides act through four distinct receptors: Pth1r, Pth2ra, Pth2rb, and Pth3r.

PTH is the primary regulator of calcium and phosphate homeostasis in humans. However, the functional diversification of the PTH family in teleosts has resulted in more complex regulatory mechanisms, particularly regarding mineral balance and skeletal development. The co-orthologs of PTH in teleosts, *pth1a* and *pth1b*, have not yet been clearly linked to bone mineral metabolism. Indeed, the extant literature on this topic is somewhat conflicting, with studies by Kwong & Perry (2015), Suzuki et al. (2011), Canario et al., (2006), and Rotllant et al., (2005) all reporting divergent functions. Furthermore, the *pth2* gene, the teleost ortholog of the human PTH2 (formerly known as *tip39*), appears to be primarily involved in regulating anxiety and nociception, with no clear role in mineral homeostasis identified to date (Anneser et al., 2022; Dobolyi et al., 2010). Conversely, the *pth3a* (*pthlha*) and *pth3b* (*pthlhb*) co-orthologs are expressed in multiple tissues and do appear to play a role in mineral homeostasis acting through endocrine and local pathways (Abbink & Flik, 2007). The most recently identified member of the PTH family, *pth4*, is expressed in two clusters of neurons in the lateral hypothalamus and has been shown to affect mineral homeostasis (Suarez-Bregua, Torres-Nuñez, et al., 2017). Notably, ectopic expression of *pth4* driven by *eef1a1* results in impaired mineralization and altered expression of key phosphate (Pi) homeostasis genes, including *fgf23, phex, entpd5* or *slc34a1a* (Suarez-Bregua, Torres-Nuñez, et al., 2017).

Phosphate plays a critical role in skeletal development, and teleosts have evolved specific regulatory mechanisms to maintain phosphate homeostasis. Unlike calcium, which is absorbed directly from the water through the gills and skin, fish primarily obtain phosphate from their diet (Lall & Kaushik, 2021). Following absorption in the intestine, phosphate is stored alongside calcium as hydroxyapatite crystals in bone and scales, which acts as major reservoirs. The regulation of phosphate release and deposition in the bone matrix is mediated by osteoblasts and osteoclasts, influenced by endocrine signals, including hormones such as Pth1a, Pth3a (Pthlha), and potentially Pth4. These hormones regulate phosphate homeostasis by acting on various tissues, particularly the kidney, which plays a crucial role in phosphate reabsorption and excretion.

The present study aims to investigate the function of Pth4 in zebrafish, with a particular focus on its role within the broader regulatory networks of mineral homeostasis. Utilizing CRISPR/Cas9 technology, we have generated *pth4* knockout zebrafish and assessed the phenotypic effects through the utilization of light-sheet fluorescence microscopy. Furthermore, calcium/phosphate (Ca/P) ratios in scales were measured to explore changes in mineral composition. Additionally, RNA sequencing (RNA-seq) was performed across six key tissues—intestine, brain, kidney, gills, scales, and bone—to examine the transcriptomic consequences of Pth4 loss.

## Materials and methods

### Zebrafish Husbandry and Ethical Statement

The present study was performed using wild-type zebrafish (Tübingen strain, Nüsslein-Volhard Lab) and *pth4* knockout zebrafish, which were reared in the facilities of the Institute of Marine Research (IIM-CSIC), Vigo, Spain. All experimental procedures were approved by the Institutional Animal Care and Use Committee of the IIM-CSIC in accordance with the Royal Decree (53/2013) and the European directive (2010/63/EU) for the protection of experimental animals (Reference: ES360570202001 / 18 / FUN. 01 / BIOL AN. 08/JRM).

### Generation of *pth4* Knockout Mutants (*pth4*^*KO*^)

The *pth4* loss-of-function mutation was induced using the CRISPR/Cas9 technology, following a protocol originally adapted from Bassett et al., (2013). Two potential target sites (TS1 and TS2) within the *pth4* gene were identified using the CHOPCHOP web tool (Labun et al., 2019) (TS1: 5’-GACTCTGAATGGAGCGACCG-3’ and TS2: 5’-GACCGCGGTCATGCATCAGC-3’). To synthesize each guide RNA (gRNA), a template-free PCR was performed using two oligonucleotides: a scaffold oligo (5′-GATCCGCACCGACTCGGTGCCACTTTTTCAAGTTGATAACGGACTAGCCTTAT TTTAACTTGCTATTTCTAGCTCTAAAAC-3′) and two different gene-specific oligos (GS1: 5’-AATTAATACGACTCACTATAGACTCTGAATGGAGCGACCGGTTTTAGAGCTAG AAATAGC-3’ and GS2: 5’-AATTAATACGACTCACTATAGACCGCGGTCATGCATCAGCGTTTTAGAGCTAG AAATAGC-3’). The PCR was carried out in a 20 μL reaction volume containing: 10 μL of 2× Phusion High-Fidelity PCR Master Mix Buffer (New England Biolabs, Ipswich, MA, USA), 1 μL of each gene-specific oligo (10 μM), 1 μL of gRNA scaffold oligo (10 μM) and nuclease-free water to a final volume of 20 μL. The PCR conditions that were utilized are outlined low: initial denaturation 98°C for 30 seconds, followed by 40 cycles of denaturation at 98°C for 10 seconds, annealing at 60°C for 10 seconds, and extension at 72°C for 15 seconds. The final extension step was performed at 72°C for 10 minutes. The resulting 125 base pair (bp) DNA fragment was purified using the DNA Clean&Concentration-5 kit (Zymo Research, Orange, CA, USA) according to the manufacturer’s instructions. The purified DNA fragment was then employed as a template for *in vitro* transcription with the MEGAscript T7 High yield transcription Kit (Ambion, Austin, TX, USA) in accordance with the manufacturer’s instructions.

In the experiment, both synthetized gRNAs were microinjected simultaneously at a final concentration of 25 ng/μL each, along with Cas9 mRNA (transcribed from the linearized pT3TS-nCas9n plasmid; Addgene, Watertown, MA, USA) at a 50 ng/μL concentration and phenol red solution (0.1%). Approximately 2 nL of this mixture was microinjected into each WT zebrafish embryo at the one-cell stage using a microscope (M165FC, Leica) and a MPPI-2 pressure injector (ASI systems). Uninjected embryos were used as controls. Following the injections, several mutations were identified, and three distinct potential nonfunctional mutations were raised as separate *pth4* knockout lines (*pth4*^*KO*^).

### Calcium and Phosphorus determination

In order to characterize the effect of the *pth4* gene knockout on bone mineral homeostasis, the calcium (Ca) and phosphorus (P) concentrations in fish scales were determined by means of inductively coupled plasma mass spectrometry (ICP-MS, Agilent 7900) in He mode using external calibration. Five *pth4*^*KO*^ fish and five WT fish (360 days post fertilization, dpf) were euthanized with a lethal dose of MS-222 (Sigma-Aldrich, St. Louis, MO, USA) and their scales were manually collected. Prior to analysis, the scales were thoroughly cleaned by sonication in ultrapure water for a period of 15 minutes. Thereafter, the scales were subjected to a digestion process that lasted for a period of 48 hours at room temperature in 1 mL of 69% HNO_3_ (Merck Suprapur). The digested samples were then diluted to a final volume of 15 mL with ultrapure water prior to analysis. The accuracy of the Ca and P measurements was verified by analyzing the reference materials 1640a (Trace elements in natural water; NIST, USA), CRM513 (Limestone; BAS, UK) and DORM5 (Fish protein; NRC, Canada), (Supplementary Data 1). For the CRM513 and DORM5, the same digestion procedure as for the scales was applied. The statistical significance of the Ca/P ratio was determined by two-tailed *t*-test, with *P*-values <0.05 considered statistically significant.

### Bone staining and Light-sheet Fluorescence Microscopy Imaging (LSFM)

Following the removal of scale, 10 adult fish (5 *pth4*^*KO*^ and 5 WT, 360 dpf) used for Ca/P estimation were fixed in 4% paraformaldehyde for 24 hours. Bone staining was then performed with Alizarin Red, following the protocol described by Gavaia P.J. et al. (2000), allowing clear visualization of calcified structures. Following staining, the fish were preserved in glycerol and visualized using a Leica M165FC microscope and a Zeiss Lightsheet7 imaging system (Zeiss, Oberkochen, Germany). Images were captured using a 5× NA 0.16 Plan-Neofluar detection objective with a zoom of 0.8. To detect the Alizarin Red fluorescent signal, the illumination laser was set to 561 nm and paired with an SBS LP 560 detection filter. The Pivot Scanner system was employed to eliminate shadowed areas within the field of view. Images were acquired at a resolution of 1.24□×□1.24□×□5.18 μm. For each specimen and region of interest, a suitable number of tiles were selected, ranging from 3×3 to 3×4 for the cranial region and 2×6 to 2×7 for the axial skeleton. Subsequent to this, tile stitching was completed using Zen Blue v3.8 software (Zeiss). Minimal adjustments to the images, including color correction and the removal of artifacts, were performed when necessary using Adobe Photoshop CC 2019 (Adobe Systems Inc., USA).

### RNA-seq experimental design and sample collection

For this study, 4 *pth4*^*KO*^ fish and 4 WT fish were used. Six tissues were sampled from each fish: the intestine, brain, kidney, gills, scales and bone (axial skeleton from the first vertebra to the last vertebra before the caudal fin was sampled). The fish were euthanized with a lethal dose of MS-222 (0.1 %, w/v in system water) (Sigma-Aldrich) prior to dissection, and tissue samples were collected from each individual, resulting in a total of 48 samples.

### RNA Isolation and Sequencing

Total RNA was isolated from dissected tissues using TRIzol (Thermo Fisher Scientific, Waltham, MA, USA) and subsequently purified with the RNeasy Mini Kit (Qiagen, Germany) in accordance with the manufacturer’s instructions. The RNA concentration was determined using a Qubit 4 fluorometer (Thermo Fisher Scientific) and the integrity was assessed using an Agilent 2100 bioanalyzer (Agilent Technologies, USA). Samples with an RNA integrity number exceeding 8 were selected for further analysis.

Strand-specific RNA libraries were then prepared and high-throughput sequencing was performed by BGI Genomics (Shenzhen, China). Libraries were sequenced on a DNBSEQ platform, generating 24.05 million paired-end reads of 100 bp each.

### Transcriptome Analysis

Raw read quality was assessed using FastQC v0.12.0 (Andrews, 2010), and adapter sequences, along with low-quality reads, were removed using SOAPnuke v2.1.9 (Chen et al., 2018). The cleaned sequences were then aligned to the Ensembl zebrafish genome assembly (Harrison et al., 2024, GRCz11) using STAR v2.7.11b (Bruchmann et al., 2013). Gene-level counts were subsequently determined with the STAR *--quantMode* option. Downstream analysis was performed using the R package *edgeR* v4.4 (Robinson et al., 2009). Low-count genes were filtered using the *filterByExpr* function, followed by normalization with the trimmed mean of M-values (TMM) method (Supplementary Data 2). Subsequently, differential expression testing was conducted using edgeR’s quasi-likelihood ratio test, and genes were considered differentially expressed (DEGs) if they met the criteria of FDR < 0.05 and |log□FC| > 1. Volcano plots were generated using ggplot2 v3.5.1 (Wickham, 2016) and the Upset plot was created with the ComplexHeatmap R package v2.20.0 (Gu et al., 2016). Gene ontology (GO) enrichment analysis was performed with the clusterProfiler R package v4.12.1 (Wu et al., 2021).

## Results

### Generation of *pth4* Knockout Zebrafish Lines

In order to elucidate the function of Pth4 in bone development and mineral homeostasis, a *pth4* knockout zebrafish line was generated using CRISPR/Cas9 technology. Two target sites within exon 2 of the *pth4* gene were selected, resulting in the deletion of the PTH-related domain that is essential for ligand-receptor binding (Fig. 1A). Three distinct frameshift mutations were identified: M1, which involves a 4-bp deletion and a 9-bp insertion; M2, a 17-bp deletion; and M3, a 13-bp deletion (Fig. 1B). All three mutations result in the introduction of a premature stop codon and predicted non-functional protein (Fig. 1C). The M1-derived protein consists of 50 amino acids, sharing 43 amino acids with WT Pth4; M2 yields a 53-amino acid protein with 40 shared; and M3 mutation encodes a 44-amino acid protein, sharing 41 amino acids with WT. Stable homozygous mutant lines were established and designated as follows: *pth4*^*iim20*^, *pth4*^*iim21*^ and *pth4*^*ii22*^ resulting from M1, M2 and M3 mutations, respectively. Given the absence of any discernible phenotypic difference between the three mutant lines, the present study was conducted on the *pth4*^*iim21*^ line, henceforth designated as *pth4*^*KO*^.

**Figure 1.**
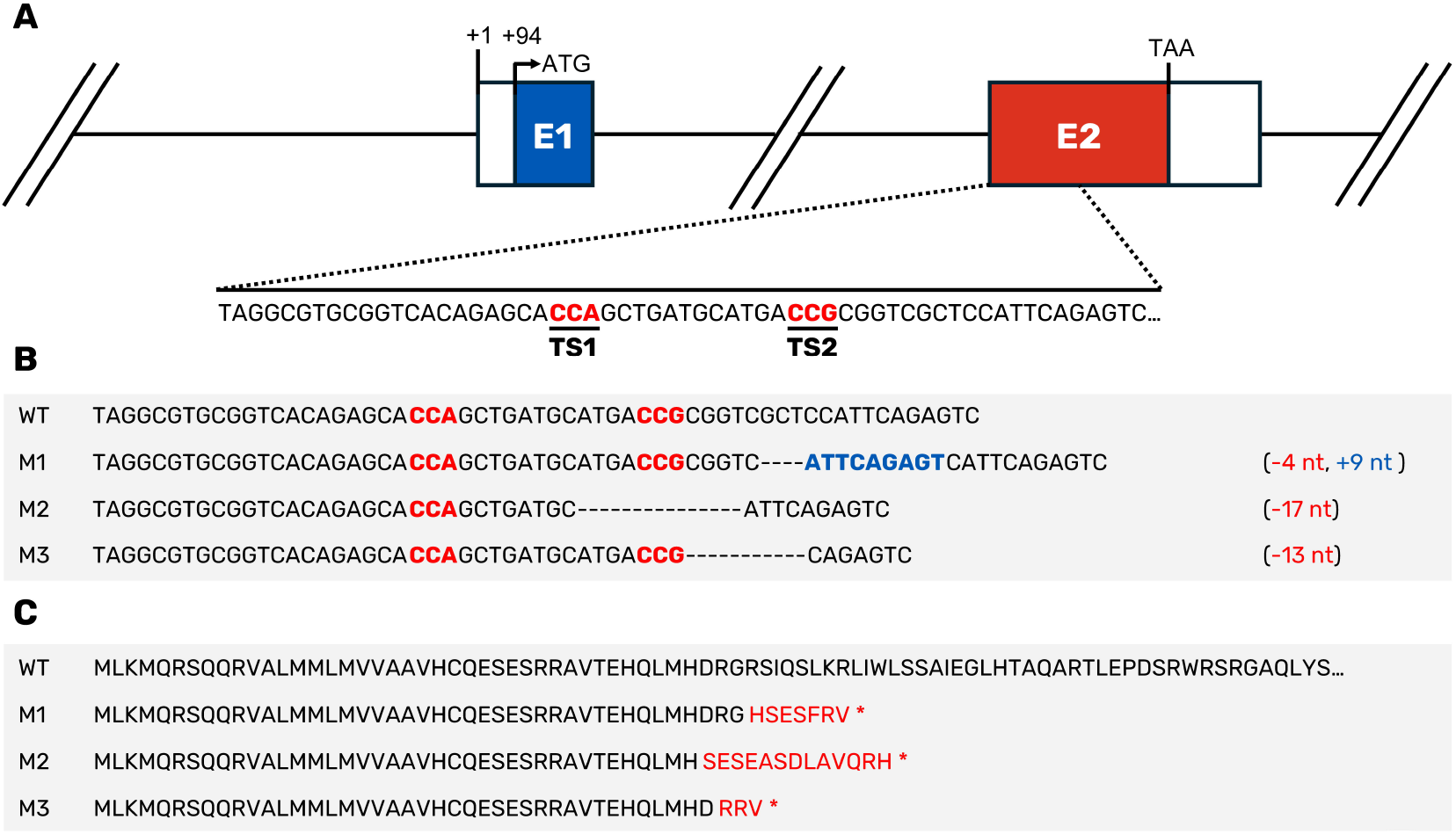
CRISPR/Cas9-induced mutations at the zebrafish *pth4* locus. (**A**) Graphical representation of the *pth4* gene. Coding exons are represented with colored boxes and the 5’-UTR and 3’UTR are represented as white boxes. (**B**) Nucleotide sequence of wild-type *pth4* (WT) and induced mutations (M1, M2 and M3). The protospacer-adjacent motif (PAM) is indicated in red, denoting the mutation target sites (TS1 and TS2). Hyphens represent deletions and nucleotides in blue represent insertions. (**C**) Predicted amino acid sequences of wild-type *pth4* and induced mutations. Amino acids in red differ from the WT sequence and asterisks indicate stop codons.

### Characterization of *pth4* Knockout Zebrafish

Alizarin red staining, which selectively binds to calcium deposits and is commonly used to visualize mineralized bone, was employed to characterize the skeletal phenotype of both larvae (14 dpf) and adult (360 dpf) *pth4*^*KO*^ and WT zebrafish. The selection of these time points enable the evaluation of both the early stages of skeletal development and the subsequent long-term maintenance of bone homeostasis and remodeling. No discernible skeletal phenotypic differences were observed between *pth4*^*KO*^ and WT larvae. However, a marked disparity in skeletal deformities was evident between adult *pth4*^*KO*^ fish and WT (Fig. 2A and B), with the former exhibiting malformed vertebrae and spinal abnormalities, including lordosis and scoliosis. These findings suggest that Pth4 plays a critical role in the maintenance of the axial skeleton during adulthood but may be dispensable during early skeletal patterning. The deformities were observed throughout the trunk and tail, with no evident positional bias (Fig. 2C, D, E, G), and were present in approximately 85% of the cases. Despite the severe vertebral deformities observed in adults, mineralization of craniofacial and appendicular bones remained unaffected (Fig. 2F, H), suggesting region-specific susceptibility to Pth4 loss.

**Figure 2.**
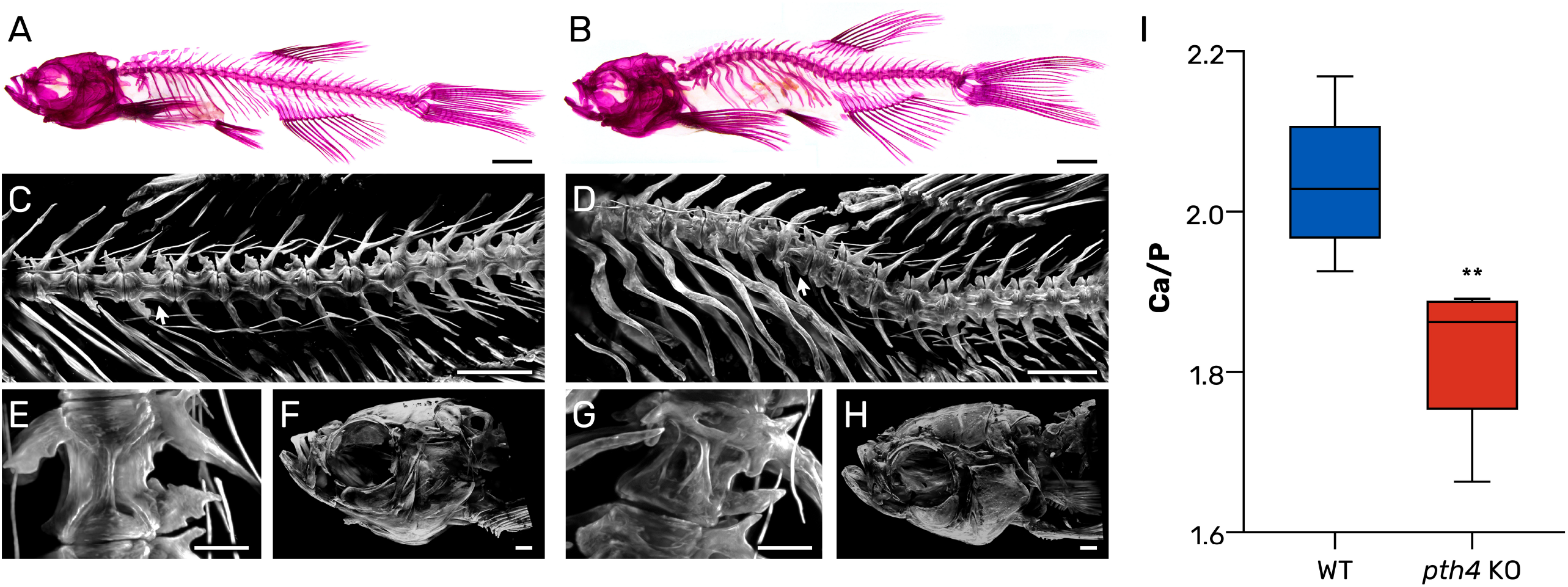
Mutant *pth4*^*KO*^ zebrafish exhibit early-onset physical decline, spinal deformities and altered Ca/P ratio in scales. (**A-B**) Lateral whole-body images of alizarin red stained WT (**A**) and *pth4*^*KO*^(**B**) adult zebrafish (360 dpf). The mutant fish display prominent deformities including lordosis, kyphosis and malformed vertebrae. (**C-D**) Light-sheet fluorescence microscopy (LSFM) images of the axial skeleton of WT (**C**) and *pth4KO* (**D**) zebrafish showing vertebrae from c8-c6 to c19. White arrows indicate the specific vertebrae highlighted in panels E and F. (**E-F**) LSFM images of vertebra c11 in WT (**E**) and *pth4*^*KO*^ (F) zebrafish. WT zebrafish exhibit well-developed vertebral structures, with the characteristic hourglass shape of the centrum clearly visible, while the mutant exhibits significant malformations. (**G-H**) LSFM images of craniofacial skeleton of WT (**G**) and *pth4*^*KO*^ (**H**) zebrafish, in which no major abnormalities were detected. Scale bar: (**A-B**) 2mm, (**C-D**) 1mm, (**E-F**) 200μm (**G-H**) 500μm. (**I**) Boxplot representing the Ca/P ratio as determined in scales. The median value is represented by a dark horizontal line, and the whiskers represent the minimum and maximum values of the data set, ** *P*<0.01.

### Effect of Deletion of *pth4* on Calcium and Phosphorus Content

To determine the impact of the *pth4* gene knockout on calcium (Ca) and phosphorus (P) content, the Ca/P ratio in zebrafish scales was measured using inductively coupled plasma mass spectrometry (ICP-MS). The mean Ca/P mass ratio in *pth4*^*KO*^ zebrafish was 1.829 ± 0.094, while in WT zebrafish, it was 2.035 ± 0.087 (mean ± standard deviation). A t-test revealed a significant difference between the *pth4*^*KO*^ and WT groups (*P*=0.007), indicating that the absence of Pth4 has a substantial effect on the calcium-to-phosphorus balance in zebrafish scales (Fig. 2I).

### Global Overview of RNA-sequencing Data

In order to assess the transcriptomic consequences of *pth4* knockout, RNA-seq was performed on six key tissues selected for their roles in calcium and phosphate homeostasis: intestine and gills (sites of phosphate and calcium absorption); brain (involved in calcium signaling); kidney (regulation of calcium and phosphate reabsorption); scales (calcium and phosphate reservoir); and together with bone, form the scale-bone axis involved in the regulatory interface for systemic mineral balance in teleosts. A total of 48 cDNA libraries were sequenced, yielding an average of 24.05 million paired-end reads per sample. After quality filtering, an average of 93.31% of reads were successfully mapped to the Ensembl zebrafish genome assembly (GRCz11), thereby ensuring the generation of high-quality data for differential expression analysis (Supplementary Data 3). Then, an assessment was made of sample clustering and variability by constructing a principal component analysis (PCA, Supplementary Data 4) and sample distance heatmap based on global gene expression profiles (Fig. 3). These analyses revealed strong within-tissue clustering, with kidney samples showing the largest transcriptional divergence between WT and *pth4*^*KO*^ groups. This finding suggests that the kidney may be particularly affected by the loss of function of the *pth4* gene, attempting to compensate for the disruption in calcium levels by altering the expression of genes involved in calcium reabsorption or excretion. It is noteworthy that tissues with analogous calcium regulatory functions, including gill and intestine (calcium and phosphate uptake), and scales or bone (calcium and phosphate reservoir), did not exhibit similar clustering patterns. This unexpected lack of correlation suggests that the loss of Pth4 may trigger distinct tissue-specific responses rather than a coordinated regulation across calcium and phosphate-regulating tissues.

**Figure 3.**
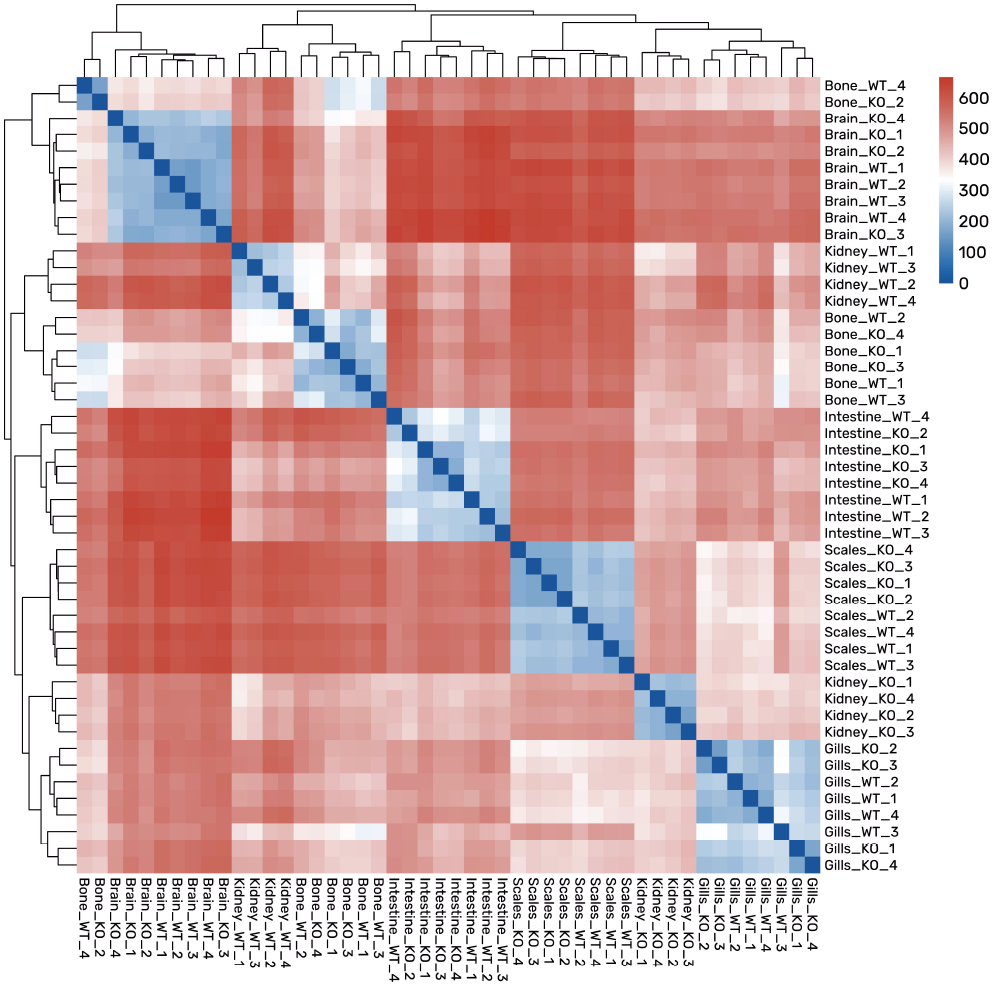
Global Overview of RNA-sequencing Data. Sample-to-sample distance heatmap. Clear within-tissue clustering is evident, with kidney samples displaying the most significant transcriptional divergence.

### Loss Of Pth4 Induces Widespread Transcriptomic Changes in Multiple Tissues

To assess tissue-specific effects of pth4 knockout, differential expression analyses were performed for each tissue by comparing WT and *pth4*^*KO*^ samples. A total of 16,902 DEGs were identified across six tissues: kidney (14,139), brain (472), intestine (510), gills (891), scales (589), and bone (323) (Supplementary Data 3, 5–10). Volcano plots were generated to visualize transcriptional changes (Fig. 4A). The kidney displayed particularly pronounced alterations, with several genes showing log_2_ fold changes exceeding 15, most of which were downregulated in *pth4* mutants. This indicates a strong transcriptional repression associated with Pth4 loss. In contrast, the remaining tissues exhibited a more balanced distribution of up- and downregulated genes.

**Figure 4.**
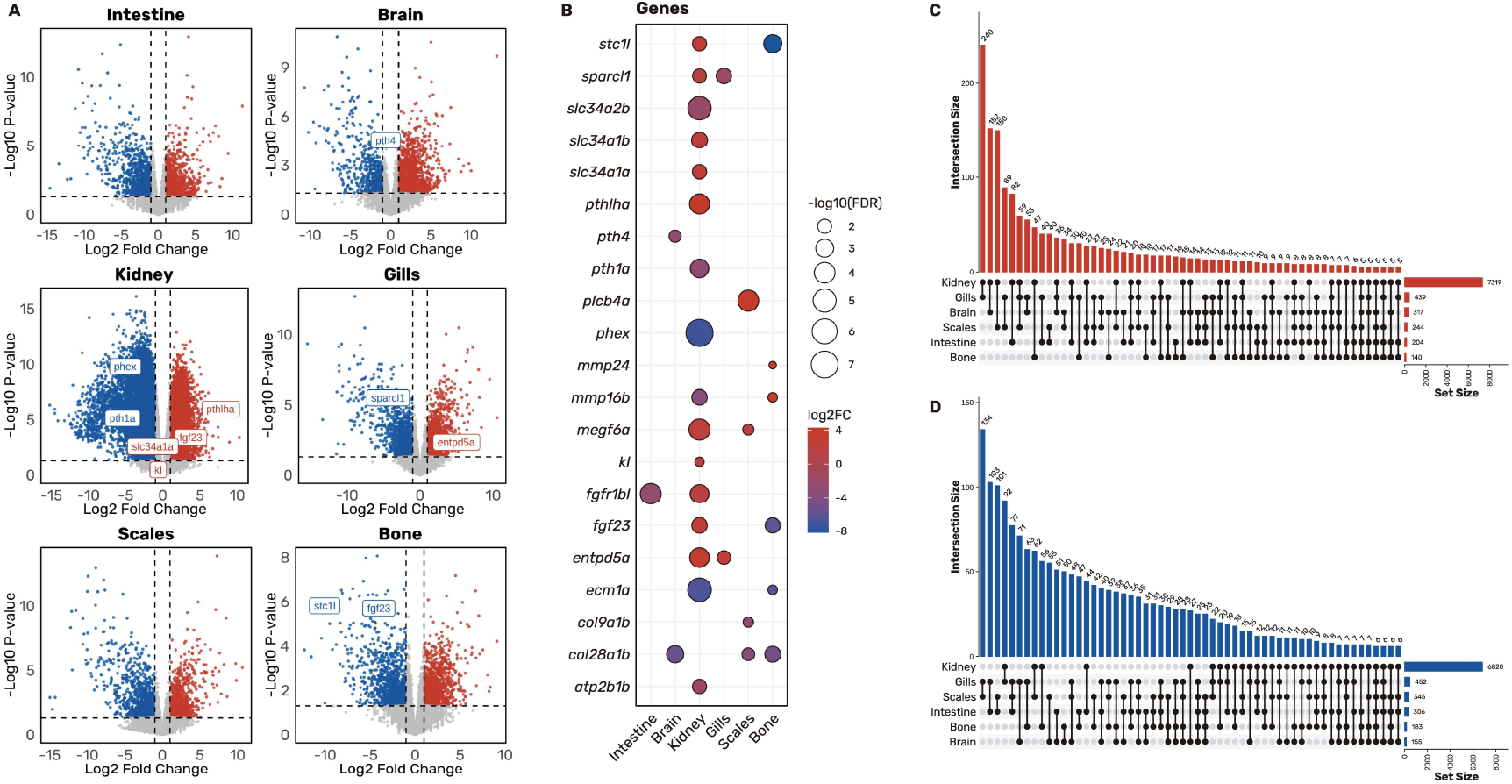
Knockout of *pth4* gene induces transcriptomic changes in mineral homeostasis-related genes in multiple tissues. (**A**) Volcano plots illustrating differentially expressed genes (DEGs) between WT and *pth4*^*KO*^ zebrafish for each tissue under study. Colored dots represent significant DEGs (p < 0.05), with red indicating upregulated genes and blue indicating downregulated genes. Horizontal and vertical dashed lines illustrate the significance thresholds (*P*<0.05 and |-Log2FC|>1, respectively). Significant genes related to mineral homeostasis are highlighted. (**B**) A Bubble plot is presented, showing selected relevant DEGs involved in mineral homeostasis across the studied tissues. The size of the bubbles indicates the statistical significance of the DEGs (-log10(FDR)), while the color indicates the log2 fold change (log2FC), ranging from blue (downregulated) to red (upregulated). (**C-D**) Upset plots showing intersections of upregulated (**C**) and downregulated (**D**) DEGs (≥5) derived from all the tissues that were studied. The total number of DEGs from each tissue is represented on the horizontal right side bar plot.

In addition to the global DEG counts, the focus was directed towards those implicated in mineral and skeletal homeostasis across all tissues (Fig. 4B). The analysis revealed that collagen genes such as *col28a1b* or *col9a1b* were downregulated in mutant fish in bone and scales, while collagen catabolic genes like *mmp24* and *mmp16b* were upregulated in bone. Furthermore, in bone tissue, there was significant downregulation of key regulators of phosphate and calcium homeostasis, including *fgf23* and the hypercalcemic hormone gene *stc1l*. Interestingly, these same genes were upregulated in the kidney, suggesting a potential compensatory response. The loss of *fgf23* and *stc1l* expression in bone may trigger their renal induction as an adaptive mechanism to preserve systemic mineral balance in the face of disrupted calcium and phosphate regulation resulting from impaired Pth4 function. This divergence of expression between tissues underscores a disruption of the bone-kidney endocrine axis. In the kidney, mineral phosphate transporter genes such as *slc34a1a* (formerly known as *npt2a*) or *slc34a1b* are upregulated, and specifically, *slc34a2b* is downregulated in *pth4*^*KO*^ fish. A similar dysregulation of phosphate and calcium homeostasis genes, including *kl, phex, entpd5a, atp2b1b* or Pth family genes such as *pth3a* and *pth1a*, was observed when comparing WT and mutant fish. In contrast, the intestine and brain exhibited no specific dysregulation of mineral homeostasis genes beyond those associated with general metabolic processes and the downregulation of *pth4* in the brain, likely attributed to nonsense-mediated decay.

Following the identification of tissue-specific transcriptional changes, we next focus on the occurrence of any common or coordinated responses occurred across multiple tissues. To this end, UpSet plots were utilized to examine the overlap among DEGs across tissues (Fig. 4C, D). The kidney emerged as a central node in these comparisons, sharing the highest number of upregulated genes with gills (240), followed by brain (152), scales (150) and intestine (82). In contrast, gills and scales exhibited the greatest overlap of downregulated genes (134), suggesting a common transcriptional repression pattern. It is noteworthy that only three genes were consistently found to be upregulated across all six tissues (*slc16a8, si:dkey-71l4*.*3*, and *mhc1zka*), while another three were commonly downregulated (*ccdc134, LOC568375*, and *rnps1*), suggesting a limited universal transcriptional response. Of particular interest is the observation that the scale-bone axis, which functions as a major calcium and phosphate reservoir, exhibited 18 shared upregulated genes in *pth4* mutant fish compared to 42 downregulated genes, suggesting a coordinated suppression of mineral storage functions. A similar phenomenon was observed in the gills and intestine, which are the primary sites for calcium and phosphate uptake, respectively. These tissues shared a high number of downregulated genes (103), compared to only 40 upregulated, indicating potential impairment of calcium and phosphate absorption mechanisms. The kidney exhibited robust transcriptional connections with multiple tissues, including gills (240 upregulated, 92 downregulated), suggesting its involvement in calcium inflow and excretion; with scales (150 up, 62 down), indicative of its involvement in mobilization cycles; and with intestine (82 up, 44 down), potentially associated with calcium and phosphate absorption/excretion balance. Furthermore, the brain also shared the most DEGs with the kidney (152 upregulated, 27 downregulated) followed by the gills (55 upregulated, 71 downregulated).

Collectively, these observations imply that the absence of Pth4 instigates a complex and tissue-specific remodeling of gene expression, with the kidney potentially functioning as a central hub in the compensatory response. Furthermore, the asymmetric overlap of DEGs, the increased repression of specific axes, and the mixed responses in axes, suggest a disruption in the fine-tuned coordination normally governing calcium and phosphate homeostasis.

### Functional Enrichment of Tissue-specific and Shared DEGs Across Multiple Tissues

To assess the functional impact of the transcriptomic alterations, a GO enrichment analysis of biological processes was performed for both upregulated and downregulated DEGs across individual tissues (Supplementary Data 11-12), as well as for shared gene sets identified in tissue intersections (Supplementary Data 13-14).

Upregulated DEGs exhibited a consistent and robust enrichment of immune-related processes across all tissues. Enriched GO terms included response to cytokines (GO:0034097), positive regulation of immune system process (GO:0002684), and regulation of immune response (GO:0050776). Additional enrichment was observed for positive regulation of ERK1 and ERK2 genes (GO:0000187) and ERK1 and ERK2 cascade (GO:0070371), suggesting widespread immune and stress-response activation in the absence of *pth4*.

Tissue intersection analysis reinforced this immune-dominant transcriptional profile. For example, kidney and gills shared upregulated genes enriched in defense response to other organism (GO:0098542), innate immune response (GO:0045087), and positive regulation of cytokine production (GO:0001819), driven by inflammatory mediators such as *ccl20l, ccl19*, and *ccl35*.*2*. Similar enrichment was observed in scales and kidney, with upregulated genes involved in inflammatory response (GO:0006954), chemokine-mediated signaling pathway (GO:0070098), and cytokine-mediated signaling pathway (GO:0019221). In brain and kidney, shared upregulated DEGs were associated with epithelial and junctional processes including adherens junction organization (GO:0034332) and cell junction assembly (GO:0034329), with increased expression of *cldnb, cldn3c*, and *cdh1*. Smaller intersections, such as between scales and bone, revealed enrichment for collagen catabolic processes (GO:0030574) and positive regulation of ERK1 and ERK2 cascade (GO:0070374), further supporting a shift in tissue remodeling dynamics.

In addition to the widespread immune-related gene activation, several intersections of downregulated genes were identified across tissues, though these gene sets were generally smaller and exhibited fewer enriched GO categories. The largest shared set of downregulated DEGs was observed between gills and scales (134 genes), though without significant enrichment, indicating a heterogeneous suppression of transcripts in these external, calcified tissues. The kidney and gills shared 77 downregulated genes, including the neurotrophic factor *fgf14*, but no GO terms reached statistical significance. By contrast, co-downregulation between the intestine and gills (103 genes) and intestine and scales (101 genes) revealed more coordinated transcriptomic changes. A core set of 77 genes jointly downregulated in gills, scales, and intestine showed significant enrichment for MAPK/ERK signaling processes, particularly positive regulation of ERK1 and ERK2 *cascade* (GO:0070374). This cluster was marked by the suppression of key growth factor signaling components such as *pdgfaa* (Platelet-Derived Growth Factor AA). In addition, several collagen-related genes exhibited coordinated downregulation in a tissue-specific manner: for example, *col17a1a* (collagen type XVII) was reduced in both kidney and scales, while *col28a1b* (collagen type XXVIII) was consistently downregulated in bone, scales, and brain.

Taken together, these findings suggest that *pth4* loss triggers a systemic immune activation coupled with tissue-restricted downregulation of extracellular matrix and MAPK/ERK signaling components. While immune-related upregulation was broadly conserved across tissues, the downregulated networks appear more specialized, potentially reflecting local adaptations to impaired mineral homeostasis or developmental signaling.

## Discussion

The parathyroid hormone (PTH) family plays a central role in calcium and phosphate homeostasis and is essential for maintaining bone integrity in vertebrates. In teleost fish, whole-genome duplications have led to the expansion of the PTH family, giving rise to peptides with specialized and sometimes divergent functions. Among these, Pth4 has emerged as a key regulator of mineral balance and skeletal integrity; however, its specific role within the PTH family remains incompletely understood (Suarez-Bregua, Torres-Nuñez, et al., 2017). The present study aims to elucidate the function of Pth4 in zebrafish by generating a *pth4* knockout model using CRISPR/Cas9 technology.

In Pth4-deficient zebrafish, we observed pronounced spinal deformities in adults along with widespread, tissue-specific transcriptomic changes, underscoring the essential role of Pth4 in adult mineral homeostasis. Notably, while larvae appeared phenotypically normal, the absence of Pth4 in adults led to the early-onset physical decline and severe skeletal abnormalities. This temporal discrepancy suggests that Pth4 is dispensable during early development but becomes critical for maintaining skeletal integrity in adulthood. This finding is consistent with previous studies, where both the ectopic expression of Pth4 and the targeted laser ablation of Pth4-expressing cells in zebrafish resulted in defective skeletal mineralization and dysregulation of osteoblast differentiation markers (Suarez-Bregua, Saxena, et al., 2017; Suarez-Bregua, Torres-Nuñez, et al., 2017).

Further supporting these phenotypic observations, ICP-MS-based elemental analysis of scales from *pth4*^*KO*^ zebrafish revealed a significant reduction in the Ca/P mass ratio compared to their wild-type counterparts. This imbalance was primarily driven by a modest increase in phosphate levels, accompanied by a slight decrease in calcium concentrations. This shift in mineral composition is particularly relevant, as the Ca/P ratio is a key determinant of hydroxyapatite crystallinity and overall bone stability. In vertebrates, the optimal Ca/P mass ratio in bone is approximately 2.15, a value that closely aligns with that observed in WT zebrafish in this study (Bonjour, 2011; Nikiforidou et al., 2020). In contrast, the lower Ca/P ratio observed in *pth4*^*KO*^ animals derived in this case from a relative excess of phosphate and/or deficiency of calcium, conditions known to compromise the quality and strength of mineralized structures (Hou et al., 2022).

Altered Ca/P ratios have been associated with decreased bone mineral density and increased fracture risk in various vertebrate models (Costa et al., 2018; Huttunen et al., 2007). This biochemical imbalance correlates directly with the skeletal deformities we observed in adult *pth4*^*KO*^ zebrafish. The absence of Pth4 most likely disrupts calcium-phosphate homeostasis, leading to impaired bone mineralization and weakened skeletal integrity, rendering structures such as the spine more vulnerable to deformities, including lordosis. These findings are in line with the established roles of Pth4 and other members of PTH family in regulating calcium and phosphate balance across vertebrates species (Guerreiro et al., 2007; Suarez-Bregua, Cal, et al., 2017).

To understand the transcriptomic impact of the Pth4 loss, an RNA-seq analysis was performed across six tissues. This revealed that the effects of Pth4 loss are widespread but highly tissue-specific. Among all the tissues that were studied, the kidney, a central organ in phosphate reabsorption and excretion, showed the most pronounced transcriptional changes. In zebrafish, the kidney plays a crucial role in regulating phosphate levels, primarily through sodium-phosphate co-transporters like *slc34a1a* (formerly *npt2a*) and *slc34a1b*, which facilitate phosphate reabsorption from the glomerular filtrate. The present study observed upregulation of *slc34a1a* and *slc34a1b*, alongside downregulation of *slc34a2b* in *pth4*^*KO*^ zebrafish, suggesting a complex response to phosphate handling. The distribution of these transporters along different segments of the nephron is also of interest. While there is a limited number of specific studies on the precise localization of *slc34a1a* and *slc34a1b* in zebrafish, in mammals, the orthologue *SLC34A1* is mainly localized in the proximal tubules, where it is responsible for bulk phosphate reabsorption (Wagner et al., 2014). In zebrafish, *slc34a2b* has been reported to have a more distal localization and plays a complementary role in phosphate regulation, particularly in fine-tuning phosphate reabsorption (Verri & Werner, 2019). This dysregulation in the expression of phosphate transporters could be indicative of a compensatory mechanism or a maladaptive response aimed at maintaining phosphate balance in the face of disrupted homeostasis caused by Pth4 deficiency. The altered expression of these transporters across nephron segments is likely to contribute to the observed disruption in mineral balance, particularly the reduction in the Ca/P ratio and the associated skeletal deformities in *pth4*^*KO*^ zebrafish. Another important dysregulated gene found in the kidney is the fibroblast growth factor 23 (*fgf23*), which promotes phosphate excretion by regulating renal phosphate transporters. In zebrafish, *fgf23* originates primarily from the corpuscles of Stannius (CoS), small endocrine structures located in direct contact or embedded within the kidney (Klingbeil et al., 2022). In *pth4*^*KO*^ fish, we observed upregulation of *fgf23* in the kidney, suggesting that increased CoS activity may be a response to phosphate imbalance. This was accompanied by a downregulation of *phex*, a phosphate-regulating endopeptidase, which may destabilize *fgf23* control despite functional differences from mammalian systems (Mangos et al., 2012). Additionally, the Fgf23 co-receptor klotho (*kl*) was found to be overexpressed in the kidney. The klotho co-receptor has been demonstrated to bind to Fgf receptors, which may result in an increase in Fgf23 signaling (Mangos et al., 2012; Singh et al., 2019). A similar pattern of dysregulation was observed for *stc1l*, that acts as a hypocalcemic factor that promotes renal phosphate reabsorption. Like *fgf23, stc1l* was upregulated in the kidney of *pht4*^*KO*^ fish. The increased expression of both genes in the kidney is likely aims to correct the systemic mineral imbalance by enhancing phosphate excretion (via Fgf23) and modulating calcium regulation (via Stc1l). This dual response provides a plausible explanation for the altered Ca/P ratio observed in *pth4*^*KO*^ fish and the resulting skeletal abnormalities in adult fish.

Looking at the skeletal tissues analyzed (scales and bone), collagen-related genes such as *col28a1b* and *col9a1b* were found to be downregulated, whereas matrix metalloproteinases, such as mmp*13, mmp24*, and *mmp16b* were upregulated in bone. This indicates an increased collagen degradation process. This imbalance between collagen production and degradation compromises the integrity of the bone matrix, further exacerbating the structural defects observed in *pth4*^*KO*^ zebrafish. The degradation of the extracellular matrix by metalloproteinases contributes to the deterioration of bone tissue and the reduction of its mechanical strength, ultimately resulting in conditions such as lordosis-kyphosis and malformed vertebrae (Cotti et al., 2024; Dietrich et al., 2021). Furthermore, extensive transcriptional changes have been observed in other mineral-regulating tissues, such as gills and scales. Conversely, the gut and brain exhibited a lesser degree of DEGs. The brain exhibited predominantly diminished *pth4* expression, which is likely indicative of transcriptional decay due to nonsense-mediated decay.

In addition to the classical mineral regulators, the transcriptomic data also revealed significant changes in immune and stress response genes in *pth4*^*KO*^ zebrafish. The upregulation of immune-related genes is likely to represent an indirect response to skeletal stress and inflammation, rather than a direct effect of *pth4* loss. The impaired bone mineralization and weakened skeletal integrity that characterize this condition can activate inflammatory signaling, which in turn promotes osteoclastogenesis (the differentiation and activation of bone-resorbing osteoclasts). This initiates a self-amplifying loop, where inflammation drives further bone degradation, exacerbating skeletal damage (Bergen et al., 2019; Deng et al., 2021; Terashima & Takayanagi, 2018). In support of this theory, we have observed widespread upregulation of inflammatory mediators such as *ccl20l, ccl19*, and *ccl35*.*2*, particularly in the kidney, scales, and gills. GO enrichment consistently highlighted immune-related biological processes, including the cytokine-mediated signaling pathway, chemokine activity, and innate immune response. These immune alterations were accompanied by enrichment of the ERK1/ERK2 cascade and the MAPK signaling pathway. Consequently, the MAPK pathway components exhibited dysregulation in the kidney, with *map3k1* and *jun* displaying upregulation and *mapk1* and *mapk3* displaying downregulation. In skeletal tissues, chronic inflammation is known to activate MAPK signaling, which in turn promotes the expression of matrix metalloproteinases involved in the breakdown of the extracellular matrix. The significant upregulation of *mmp13, mmp24*, and *mmp16b* in bone tissue may thus be partly driven by this inflammatory–MAPK axis, contributing to vertebral malformations and reduced skeletal strength. Collectively, these findings lend to support a model in which *pth4* deficiency triggers immune and stress responses across multiple tissues, thereby amplifying mineral imbalance and bone pathology through inflammatory signaling cascades.

In light of previous findings that Pth4 binds to the Pth1r receptor (Suarez-Bregua, Torres-Nuñez, et al., 2017), it was initially hypothesized that *pth4* knockout would result in the downregulation of Pth1r-associated signaling pathways. Surprisingly, our data revealed a marked upregulation of key downstream components, such as the ERK1/2 signaling cascade and metalloproteinase genes, including *mmp13*. This suggests that the loss of Pth4 may trigger compensatory mechanisms involving other PTH family members. Among these, *pth1a*, the main ortholog of mammalian PTH, was found to be downregulated in *pth4*^*KO*^ tissues, consistent with an impaired systemic regulation of calcium and phosphate. In contrast, *pth3a*, the zebrafish equivalent of PTH-related protein (PTHrP), was significantly upregulated, potentially reflecting a compensatory activation to mitigate skeletal and mineral homeostasis disturbances. While *pth1a* primarily mediates endocrine regulation through *pth1r, pth3a* acts more locally in bone and cartilage, promoting chondrocyte proliferation and preventing premature differentiation (Bergen et al., 2019; Guerreiro et al., 2007). The upregulation of *pth3a* may represent an attempt to preserve skeletal integrity in the absence of *pth4*, by maintaining local paracrine signaling crucial to matrix remodeling and development. These opposing expression patterns highlight the functional divergence within the PTH family and underscore the tissue-specific redundancy of these peptides in zebrafish. Consistent with observations in mammals, the PTH-FGF23 axis appears to remain functionally conserved: *pth4* loss triggered aberrant *fgf23* expression in kidney and bone, altered *klotho* and *phex* expression, and disrupted phosphate transporters, ultimately disturbing mineral homeostasis (Deng et al., 2021; Lanske & Razzaque, 2014; Quarles, 2003). The involvement of multiple PTH family members in this regulation emphasizes the redundancy and complexity of these pathways, with different peptides potentially compensating for one another depending on the tissue context or developmental stage.

In conclusion, this study establishes Pth4 as a key regulator of skeletal integrity and mineral homeostasis in zebrafish. The loss of *pth4* resulted in axial skeletal deformities, a decreased Ca/P ratio in scales, and broad transcriptomic alterations across multiple tissues. Of particular note were the dysregulation of phosphate transporters, the activation of the Fgf23/Klotho signaling axis, and the upregulation of immune and stress-responsive pathways, particularly involving ERK/MAPK signaling. These findings emphasize a systemic role for Pth4 in phosphate regulation and underscore the complex interplay between PTH family members, mineral metabolism, and skeletal maintenance. Overall, the present findings provide a valuable framework for future investigations into the PTH–FGF23 axis and the endocrine regulation of bone health in vertebrates.

## Supporting information

Description of Additional Supplementary Files

## Acknowledgements

The authors thank the Optical Microscopy & Image Analysis Service at IIM-CSIC, particularly Laura Gómez-Montoro and Lucía Sánchez-Ruiloba, for their assistance with light-sheet fluorescence microscopy and Susana Otero for maintenance of the experimental animals and assistance during sampling.

## Author contributions

Conceptualization, P.S.-B. and J.R.; Methodology, L.M.-M., P.S.-B., L.G.-P., C.C.-P. and A.C.G; Validation, L.M.-M., L.G.-P., E.B.-D and C.C.-P.; Formal analysis, L.M.-M.; Resources, J.R.; Data curation, L.M.-M..; Writing—original draft preparation, L.M.-M.; Writing—review & editing, P.S.-B., L.G.-P., E.B.-D., A.C.-G. and J.R.; Supervision, J.R.; Project administration, J.R.; Funding acquisition, J.R. All authors have read and agreed to the published version of the manuscript.

## Data Availability statement

The RNA-seq data corresponding to the different tissues of WT and *pth4*^*KO*^ zebrafish has been deposited in the Gene Expression Omnibus public (GSE294090).

