## Supplementary material for "Genetic ablation of Pth4 disrupts calcium-phosphate balance, bone development, and kidney transcriptome in teleosts": Description of Additional Supplementary Files

**File Name: Supplementary Data 1**

**Description**: Results of the analysis of Ca and P in the reference materials 1640a, CRM513 and DORM5

**File Name: Supplementary Data 2**

**Description**: Matrix of normalized gene counts for each studied sample

**File Name: Supplementary Data 3**

**Description**: Table summarizing global results from RNA sequencing, mapping and identified DEGs

**File Name: Supplementary Data 4**

**Description**: Principal Component Analysis (PCA) plot illustrating the sample similarity between the 48 samples. The colour coding system employed in this study is delineated in the accompanying legend, with hues denoting specific tissue types and the corresponding genotype for each sample

**File Name: Supplementary Data 5**

**Description**: Matrix of normalized counts of total differentially expressed genes from Intestine samples

**File Name: Supplementary Data 6**

**Description**: Matrix of normalized counts of total differentially expressed genes from Brain samples

**File Name: Supplementary Data 7**

**Description**: Matrix of normalized counts of total differentially expressed genes from Kidney samples

**File Name: Supplementary Data 8**

**Description**: Matrix of normalized counts of total differentially expressed genes from Gills samples

**File Name: Supplementary Data 9**

**Description**: Matrix of normalized counts of total differentially expressed genes from Scales samples

**File Name: Supplementary Data 10**

**Description**: Matrix of normalized counts of total differentially expressed genes from Bone samples

**File Name: Supplementary Data 11**

**Description**: Combined GO-BP enrichment results for upregulated DEGs across all six tissues (Bone, Brain, Gills, Intestine, Kidney, Scales). Includes a “Tissue” column indicating the origin of each term

**File Name: Supplementary Data 12**

**Description**: Combined GO-BP enrichment results for downregulated DEGs across all six tissues (Bone, Brain, Gills, Intestine, Kidney, Scales). Includes a “Tissue” column indicating the origin of each term

**File Name: Supplementary Data 13**

**Description**: Combined GO-BP enrichment results for up- and down-regulated DEGs for each intersection of the Upset plot. Includes an “Intersection” column indicating the intersection identifier and a “Genes” column with semicolon-delimited Ensembl IDs per intersection
