## Supplementary Data 3 for "Genetic ablation of Pth4 disrupts calcium-phosphate balance, bone development, and kidney transcriptome in teleosts"

|  | **Supplementary Table 1**. Global summary of RNA sequencing, mapping and identified DEGs | | | | | | | | |  |  |  |  |  |
| --- | --- | --- | --- | --- | --- | --- | --- | --- | --- | --- | --- | --- | --- | --- |
|  | **Intestine** | | **Brain** | | **Kidney** | | **Gills** | | **Scales** | | **Bone** | | **Total** | |
|  | **WT** | **KO** | **WT** | **KO** | **WT** | **KO** | **WT** | **KO** | **WT** | **KO** | **WT** | **KO** | **WT** | **KO** |
| **Total Reads** | 24.00 | 24.10 | 24.05 | 23.80 | 24.05 | 24.05 | 24.05 | 24.03 | 24.08 | 24.05 | 24.08 | 24.10 | 144.3 | 144.125 |
| **Mapped Reads** | 22.33 | 22.58 | 22.73 | 22.55 | 21.80 | 22.28 | 22.45 | 22.45 | 22.50 | 22.45 | 22.48 | 22.55 | 134.275 | 134.85 |
| **Mapping rate (%)** | 92.98% | 93.68% | 94.45% | 94.63% | 90.68% | 92.60% | 93.33% | 93.43% | 93.55% | 93.28% | 93.48% | 93.65% | 93.07* | 93.54%* |
| **Total DEGs** | 510 | | 472 | | 14139 | | 891 | | 589 | | 323 | | 16924 | |
|  | * Average mapping rate (%) | | |  |  |  |  |  |  |  |  |  |  |  |
