## Supplementary figures and images for "Genetic ablation of Pth4 disrupts calcium-phosphate balance, bone development, and kidney transcriptome in teleosts"

### Supplementary Data 4

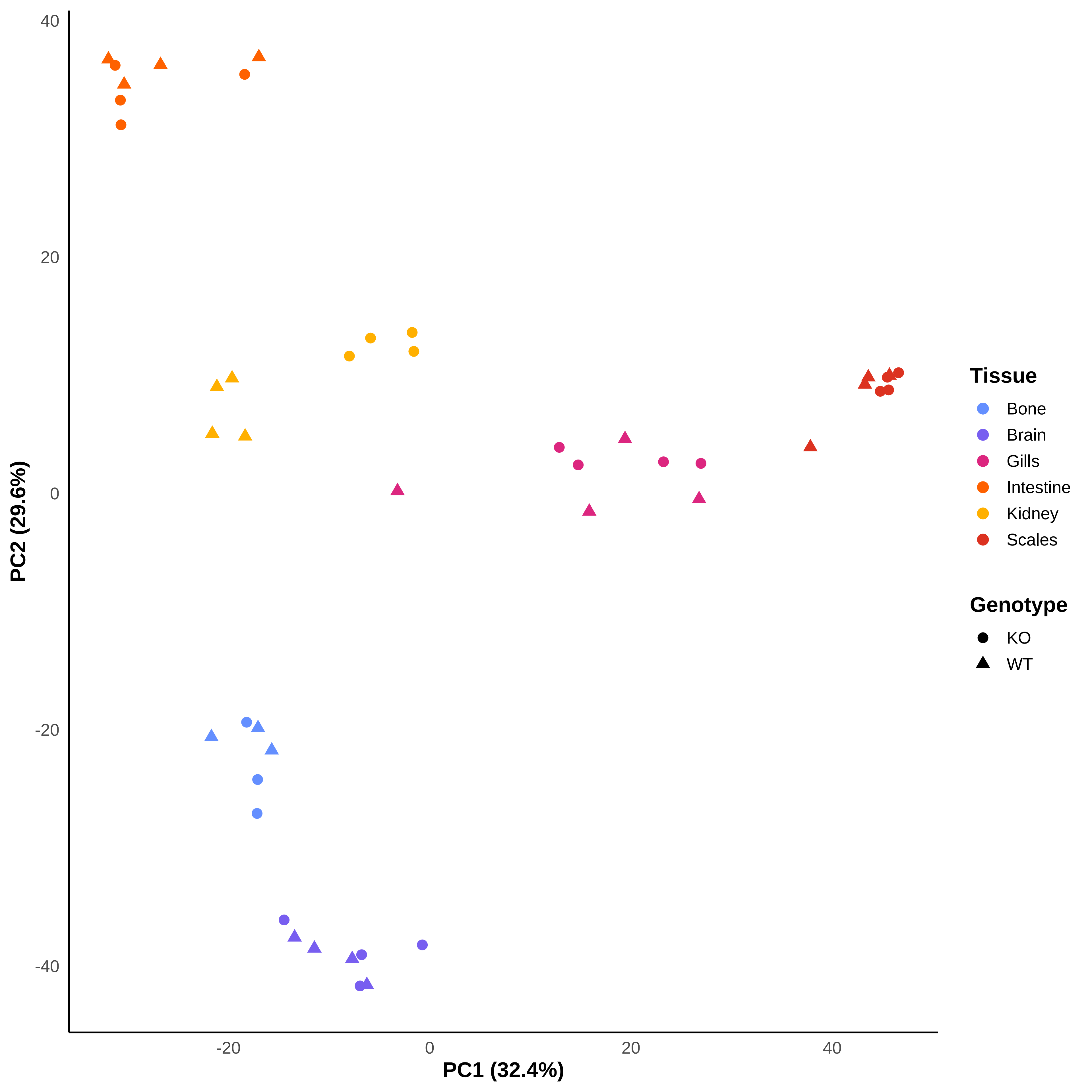
